# Optogenetic manipulation of nuclear Dorsal reveals temporal requirements and consequences for transcription

**DOI:** 10.1101/2024.11.28.623729

**Authors:** Virginia L Pimmett, James McGehee, Antonio Trullo, Maria Douaihy, Ovidiu Radulescu, Angelike Stathopoulos, Mounia Lagha

## Abstract

Morphogen gradients convey essential spatial information during tissue patterning. While both concentration and timing of morphogen exposure are crucial, how cells interpret these graded inputs remains challenging to address. We employed an optogenetic system to acutely and reversibly modulate the nuclear concentration of the morphogen Dorsal (DL), homologue of NF-κB, which orchestrates dorso-ventral patterning in the *Drosophila* embryo. By controlling DL nuclear concentration while simultaneously recording target gene outputs in real time, we identified a critical window for DL action that is required to instruct patterning, and characterized the resulting effect on spatio-temporal transcription of target genes in terms of timing, coordination, and bursting. We found that a transient decrease in nuclear DL levels at nuclear cycle 13 leads to reduced expression of the mesoderm-associated gene *snail (sna)* and partial derepression of the neurogenic ectoderm-associated target *short gastrulation* (*sog)* in ventral regions. Surprisingly, the mispatterning elicited by this transient change in DL is detectable at the level of single cell transcriptional bursting kinetics, specifically affecting long inter-burst durations. Our approach of using temporally-resolved and reversible modulation of a morphogen *in vivo*, combined with mathematical modeling, establishes a framework for understanding the stimulus-response relationships that govern embryonic patterning.

## INTRODUCTION

It is appreciated that multiple factors contribute to spatiotemporal dynamics of morphogen responses including the temporal alterations to the morphogen gradient itself, dynamics relating to signal transduction, as well as downstream interactions between target genes^2^. In particular, the duration of exposure to the morphogen has been highlighted as a crucial determinant of patterning responses, as exemplified by recent studies on Nodal, BMP, and Bicoid morphogens^3,4^. However, how the morphogen gradients are sensed, both in terms of their local concentration and the critical window they must be interpreted within to drive cell fate decision-making, remains a major question in the field. Furthermore, gene regulatory networks (GRNs) act within responding cells to interpret morphogen signals and perhaps there is built-in robustness to these systems to accommodate varied morphogen dynamics^5^. Whether morphogen spatiotemporal dynamics are central to decoding morphogen input into discrete cell fates or, instead, downstream transcriptional responses are the major drivers is a fundamental, yet unresolved, question. Distinguishing between these two alternative and potentially non-exclusive modes of action, involving direct versus indirect morphogen control, requires approaches in which the patterning process can be perturbed and studied in real time.

The dorsoventral axis of the developing *Drosophila* embryo is a well-established example of a morphogen-patterned system in which both the morphogen input and target gene outputs can be followed live in time and space^6–8^. In *Drosophila* syncytial embryos, graded input by DL is important for activating expression of target genes *snail* (*sna*) and *twist* (*twi*) in ventral regions, *ventral-neuroblasts defective* (*vnd*) in ventrolateral regions, and *short gastrulation* (*sog*) in lateral regions (Figure 1H)^9^. In addition to the spatial patterning blueprint encoded by a gradient of nuclear DL levels along the dorsoventral axis, other transcription factors (TFs) also ensure establishment of precise borders and the adoption of distinct cell fates. For instance, in ventral cells, the TFs Twist (Twi) and Snail (Sna) instruct the mesodermal fate and the subsequent epithelial-mesenchymal transition (EMT) program, essential to gastrulation^10^. Moreover, Sna directly represses the transcription of the DL target *sog* in ventral cells which form the presumptive mesoderm^11,12^. This combinatorial action of DL, Sna, and other factors such as Zelda increasingly restricts *sog* expression to lateral regions prior to gastrulation^13,14^.

**Figure 1:**
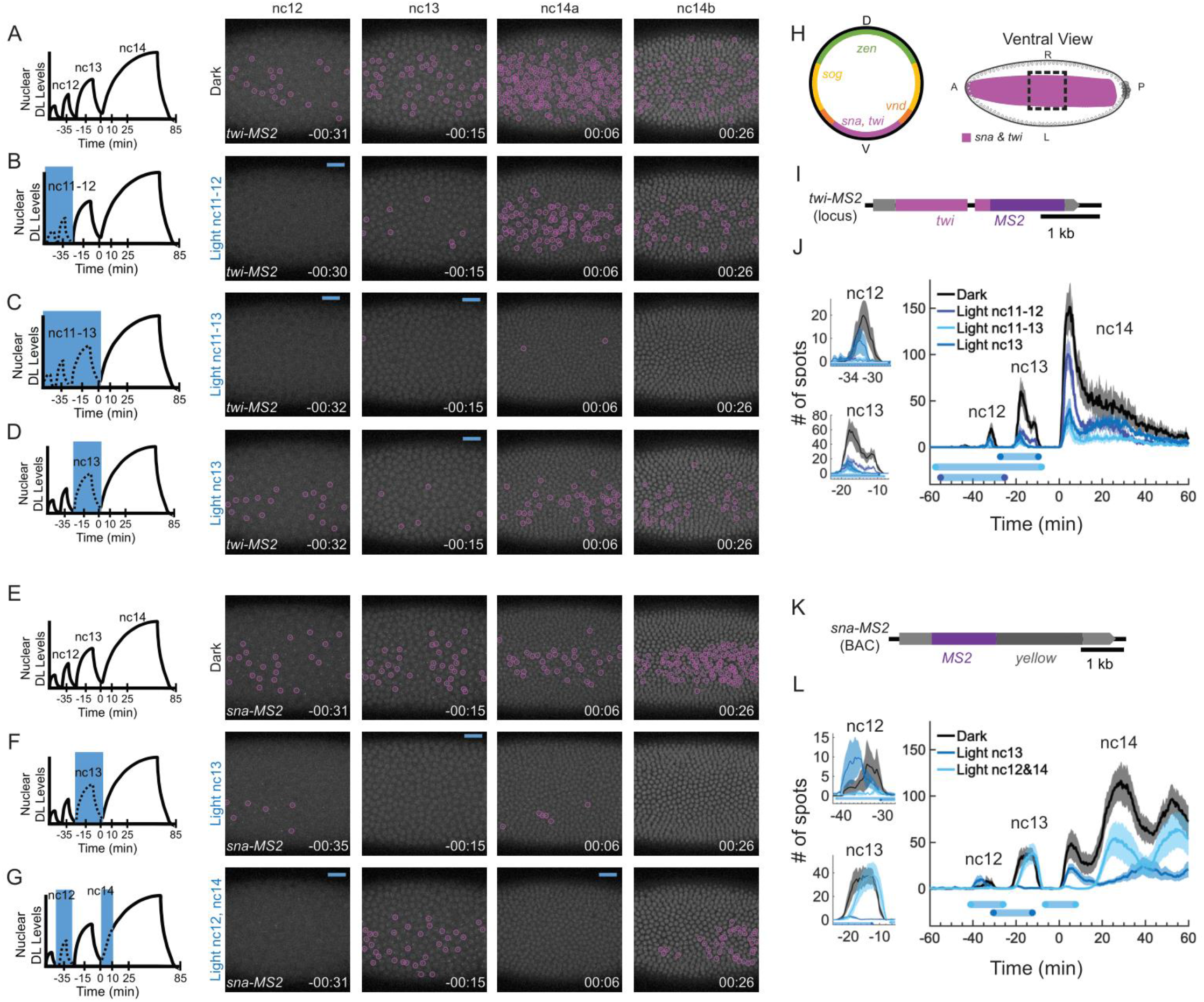
The critical window for DL activity is nc11-13 for *twi* and nc13 for *sna*. **(A-D)** *twi-MS2* (magenta) expression in the dark (A), with light at nc11-12 (B), with light at nc11-13 (C), and with light at nc13 (D). **(E-G)** *sna-MS2* (magenta) expression in the dark (E), with light at nc13 (F), and with light at nc12 and early nc14 (G). **(H)** Expression domains of DL target genes and the field of view (black dashed box) used to image *twi-MS2* and *sna-MS2* in the ventral domain. **(I**,**K)** The *twi-MS2* CRISPR construct (I) and *sna-MS2* BAC construct (K) used for tracking *twi* and *sna* expression. **(J**,**L)** Quantification of the number of active TS (mean ± SEM) detected for each time frame for *twi-MS2* (J) and *sna-MS2* (L). This metric gives the instantaneous number of active sites of transcription and is not cumulative. The black lines are in the dark (n=3 for *twi*, and n=5 for *sna*), the blue lines are with blue light at nc13 (n=3 for *twi*, and n=6 for *sna*), the darker blue line is with blue light at nc11-13 (n=4 for *twi*), and the cyan line is with blue light at nc11-12 (n=3 for *twi*) or nc12 and early nc14 (n=8 for *sna*). The blue bars represent the average illumination window for the matching condition. Nc12 and nc13 are plotted individually to the left as well as with nc14 (right). Nc14 is truncated at 60 min to ensure no embryo is gastrulating during quantification. Foci are circled in magenta for *sna* and *twi*. t=0 indicates the start of nc14. Illumination schemes are shown to the left of the image stills.

To orchestrate this complex gene regulatory network, the action of DL must be tightly regulated in both space and time. The DL gradient has been shown to be dynamic, as levels were observed to change over time^15– 17^. Although these changes in the gradient over time have been detected, the functional relevance during dorsoventral patterning remains unresolved. Recently, optogenetic tools have been used to manipulate DL levels using an opto-degron^1,18^. However, the irreversible nature of this perturbation inhibits identification of transient critical windows of DL availability^18^. By establishing a reversible optogenetic manipulation paradigm for DL nuclear levels, this study has identified gene-specific temporal requirements for DL across the dorsoventral axis. Timing is a key concept in gene regulation, particularly during development, where the dynamics of morphogens play a pivotal role in shaping gene expression that ultimately leads to patterning. Here, we have identified specific temporal requirements, showing that DL nuclear reduction in nc13 decreases *sna* expression and leads to a change in *sog* stochastic transcription properties in nc14 like decreasing the duration of a long transcriptionally inactive (OFF) state (long inter-burst periods). Our analysis further reveals changes of these transcriptional bursting properties across space, suggesting a mispatterning at the kinetic level. This analysis was only possible through a novel combination of approaches, including mathematical modeling.

## RESULTS

We used DL^LEXY^, which supports inducible DL nuclear export and is reversible (**Figure S1 A-E**)^1,19,20^, together with the MS2/MCP imaging system to monitor spatiotemporal expression of target genes *in vivo* ^21,22^. In the presence of blue light, DL^LEXY^ is translocated to the cytoplasm, and when the blue light is removed DL^mCh-LEXY^ reenters the nucleus at levels similar to before the light was applied (**Figure S2 A-D**)^1^, without significant photobleaching (**Figure S2E**). To monitor transcriptional dynamics of DL targets, we focused on four DL targets expressed in ventral, ventrolateral, or lateral regions spanning the dorsoventral axis (**Figure 1H**). We assayed gene expression of these four targets when blue light was applied during specific embryonic stages to define critical windows of DL action.

### The critical window for DL activity is nc11-13 for *twi* and nc13 for *sna*

We first examined the window of time DL must act to support target gene expression in ventral regions of the embryo, within the presumptive mesoderm. We used a published *sna*-BAC^23^ transgene and created a new *twi-MS2* allele by inserting MS2 into the endogenous *twi* locus using CRISPR/Cas9 (**Figure 1I,K**; see Methods). In the dark, *twi-MS2* expression is reliably detected from nc12 up through mid-nc14 (i.e. nc14b) but is diminished in nc14 (**Figure 1A, Movie S1**). When blue light was applied at nc11-12, *twi-MS2* signal was greatly reduced at nc12 and 13, but only moderately reduced at nc14 (**Figure 1B**). Alternatively, if embryos were exposed to light from nc11-13, *twi-MS2* signal was reduced at nc12, 13, and 14, including when returned to the dark during nc14 (**Figure 1C, Movie S1**). When illuminated at nc13, *twi*-*MS2* signal is reduced at nc13 and at nc14, but to a lesser degree than when illuminated from nc11-13 (**Figure 1D, J, Movie S1**). This indicates the DL-critical window for *twi* activation appears to be nc11-13, with nc13 appearing to be more important for the initial peak in *twi* expression than nc11-12. Importantly, the decrease in *twi* transcription under blue light is not due to photobleaching, as demonstrated by our bleaching analysis (**Figure S3A**). Taking a similar approach, we found that another ventral target gene, *sna*, exhibits a narrower critical window for DL action. When blue light is shone at nc13, *sna-MS2* signal is greatly reduced at nc13 and nc14 (**Figure 1E-G, Movie S2**). Since this reduction in the number of *sna* transcription sites (TS) was so great (**Figure 1L**), illuminating embryos from nc11-13 was not necessary for *sna*. Thus, reduction of DL levels in this short window of ∼15-20 minutes (nc13) is sufficient to silence *sna* transcription in nc13 and drastically reduce expression at following stages. We also observed that the occurrence of gastrulation is reduced in embryos illuminated at nc13 (**Figure 2A-D**). We believe that these gastrulation defects are due to the absence of Sna protein, elicited by DL nuclear reduction in nc13. We confirmed the absence of Sna protein with a more sustained light exposure of DL^LEXY^ embryos with a blue-light box set-up (**Figure S1F**; see Materials and methods). Therefore, while both *twi* and *sna* target genes are expressed in ventral regions where high levels of nuclear DL are present, these genes exhibit different temporal dependencies, with DL needing to act earlier to support *twi* and only later to support *sna*.

**Figure 2:**
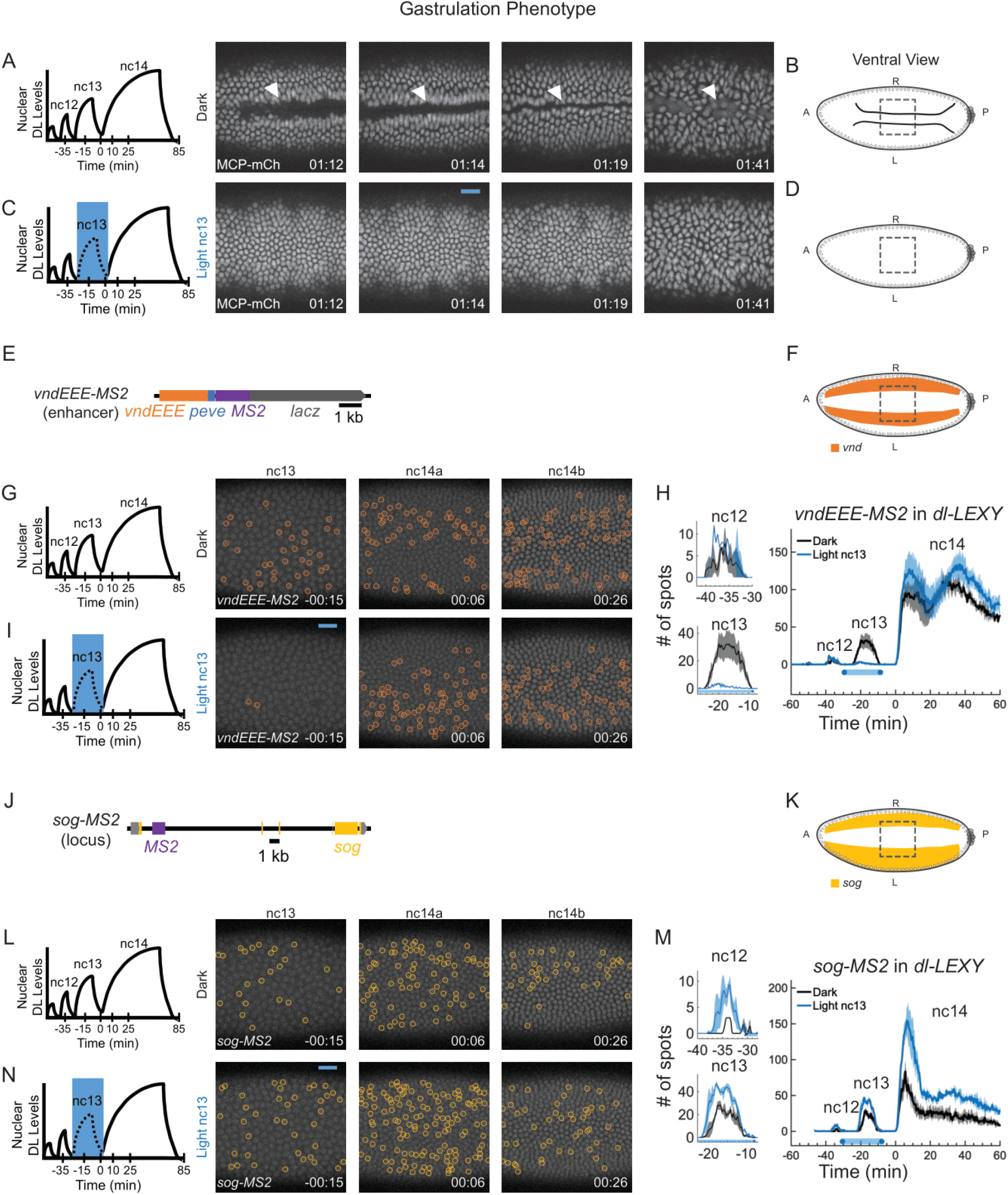
*vnd* and *sog* are retained at nc14 despite reduced nuclear DL levels at nc13. **(A**,**C)** *dl-LEXY; MCP-mCherry/TM3* was imaged for the MCP-mCherry signal with no MS2 crossed in to determine what effect the light had on gastrulation. Embryos in the dark gastrulate (A, n = 3 embryos), while embryos with light shone at nc13 fail to gastrulate completely (C, n = 3 of 4 embryos). **(B**,**D)** Schematic representing gastrulation or a failure to gastrulate, and the field of view (dashed box). **(E**,**J)** The *vndEEE-MS2* reporter construct (E) and the *sog-MS2* CRISPR construct (J) used for tracking *vnd* and *sog* expression. **(F**,**K)** Schematic of the field of view used to image ventrally for *vnd* (F) or *sog* (K). **(G**,**I**,**L**,**N)** Expression in the dark at nc13 and nc14 for *vndEEE-MS2* (G) and *sog-MS2* (L) or when illuminated with blue light at nc13 for *vndEEE-MS2* (I) and *sog-MS2* (L). Foci are circled in orange for *vnd* and yellow for *sog*. **(H**,**M)** Quantification of the number of active TS detected for *vndEEE-MS2* (H) or *sog-MS2* (M). The black lines are in the dark and the blue lines are with blue light at stages indicated (mean ± SEM; n=3 embryos for each). Nc12 and nc13 are plotted individually to the left as well as with nc14 (right). Nc14 is truncated at 60 min to ensure no embryo is gastrulating during quantification. Foci are circled in orange for *vnd* and yellow for *sog*. White arrowheads indicate the location of gastrulation. t=0 indicates the start of nc14 in (A,C,G,I,L,N).

Next, we asked whether the time of DL action or the duration of this input signal is the critical factor driving *sna* expression. To discriminate between these two alternatives, DL was exported with blue light in nc12 as well as in early nc14 within the same embryo, for a total duration of 26.3 ± 1.0 min (mean ± SEM), ∼6 min longer than the length of time blue light was given during a complete nc13 cycle duration, which was 20.5 ± 0.6 min (mean ± SEM). When we illuminated embryos with blue light during nc12 and a second time at early nc14, *sna* expression was able to recover during nc13 and late nc14 (**Figure 1G,L; Movie S2**). A similar experiment could not be performed for *twi*, as *twi* expression is difficult to detect at mid to late nc14 and recovery of *twi* expression would not be observable after removal of blue light. These results support the hypothesis that the timing of DL input in nc13 is critical for *sna* expression.

### *vnd* and *sog* persist during nc14 despite reduced nuclear DL levels at nc13

Having defined the critical temporal window required to activate ventral targets, we next questioned whether DL’s action at nc13 was also important for ventrolateral and lateral targets. To this end, we imaged *vnd*, a DL target expressed in the ventrolateral domain, and *sog*, which is expressed in the lateral domain (**Figure 1H**). We employed a reporter transgene to follow *vnd* expression in early embryos based on output of a single enhancer (vndNEE-MS2)^24^ (**Figure 2E**) and used published fly stocks with MS2 engineered into the *sog* locus (sog-MS2)^1,25^ (**Figure 2J**).

When assayed using DL^LEXY^ with illumination on the ventral side **(Figure 2F,K)**, we observed that *vnd* and *sog* responded differently to removal of Dorsal. *vndEEE* expression was acutely decreased in the presence of blue light when it was applied at either nc13 or parts of nc14, but there was no lasting effect as observed for *sna* **(Figures 2G,H,I, and S4)**, regardless of whether *vndEEE* was imaged ventrolaterally or ventrally **(Figures 2F and S4B; Movie S3)**. We quantified the number of active *vndEEE* nuclei at nc14 and found it appears largely unaffected by removal of DL at nc13 (**Figure 2H**). A previous study showed that no matter how long the blue light is shone on DL^LEXY^ embryos, *sog* expression is retained (e.g. for the entirety of nc13-14)^1^. Even though DL^LEXY^ is predominantly cytoplasmic under blue light, there is some residual, low concentration DL in the nucleus^1^ (**Figure S2**). This small amount of transient DL may be sufficient to support *sog* transcriptional activation. When embryos are illuminated with blue light on the ventral side, some *sog* was detected in regions where it should be repressed (**Figure 2K,L,N; Movie S3**). The number of *sog* active TS was higher with light at nc13 when compared to the dark condition (**Figure 2M**). Since we previously did not observe a change in *sog* expression with removal of DL^1^ and the number of *sog* TS increased instead of decreasing, this suggests a loss of repression under blue light in ventral regions.

These experiments were able to define distinct temporal requirements for DL in supporting the activation of two target genes. DL action is required early at nc11-13 for *twi*, and at nc13 for *sna*. If DL levels are reduced during these time windows, then the respective genes expression is reduced at nc14. In contrast, nc13 was not critical for activation of the ventrolateral and lateral DL target genes *vnd* and *sog*. For nc13 and 14, DL input to *vndEEE* is acutely required, but there is no long-term memory effect and gene expression recovers once DL is available again. The number of nuclei exhibiting an active *sog* TS was apparently refractory to a reduction of DL levels, even during blue light illumination. In fact, reducing DL levels on the ventral side led to an increase in *sog* TS, likely due to a loss of Sna (**Figure S1F**).

### Reduction of DL levels in nc13 does not affect the kinetics of *sog* transcription at nc13

While this quantification suggests that the probability to activate the transcription of the *sog* promoter is insensitive to blue light-induced DL nuclear export, it cannot determine if a more subtle effect may operate at the level of transcriptional kinetics. For instance, it is well appreciated that most genes are transcribed in a discontinuous manner with bursts occurring at various timescales^26^. Like other developmental genes, endogenous *sog* also exhibits a bursty transcription in the early embryo (**Movie S3; Figure 3F,H**)^25,27^. Analysis of live transcription signals shows that transcription activity is intermittent, alternating between active and several inactive periods. Mathematical models can capture this process, termed ‘bursting’, and characterize both the probability and durations of the active and inactive transcriptional periods. In addition, as *sog* is the only target gene that remains on during blue light, it was the best candidate for this analysis. Therefore, we further investigated the bursting behavior of *sog* when nuclear DL^LEXY^ levels are perturbed by blue-light at nc13, with the understanding that this would lead to a substantial reduction in *sna* expression but only a partial reduction in *twi*. To examine *sog* bursting kinetics, we imaged its transcription with high temporal resolution. Such a fast imaging set-up inevitably leads to bleaching when using a red fluorescent detector, such as the MCP-RFPT employed here. However, our bleaching study demonstrates that prior to 20 min into nc14, less than 20% of bleaching is observed, allowing adequate detection (**Figure S3B**).

**Figure 3:**
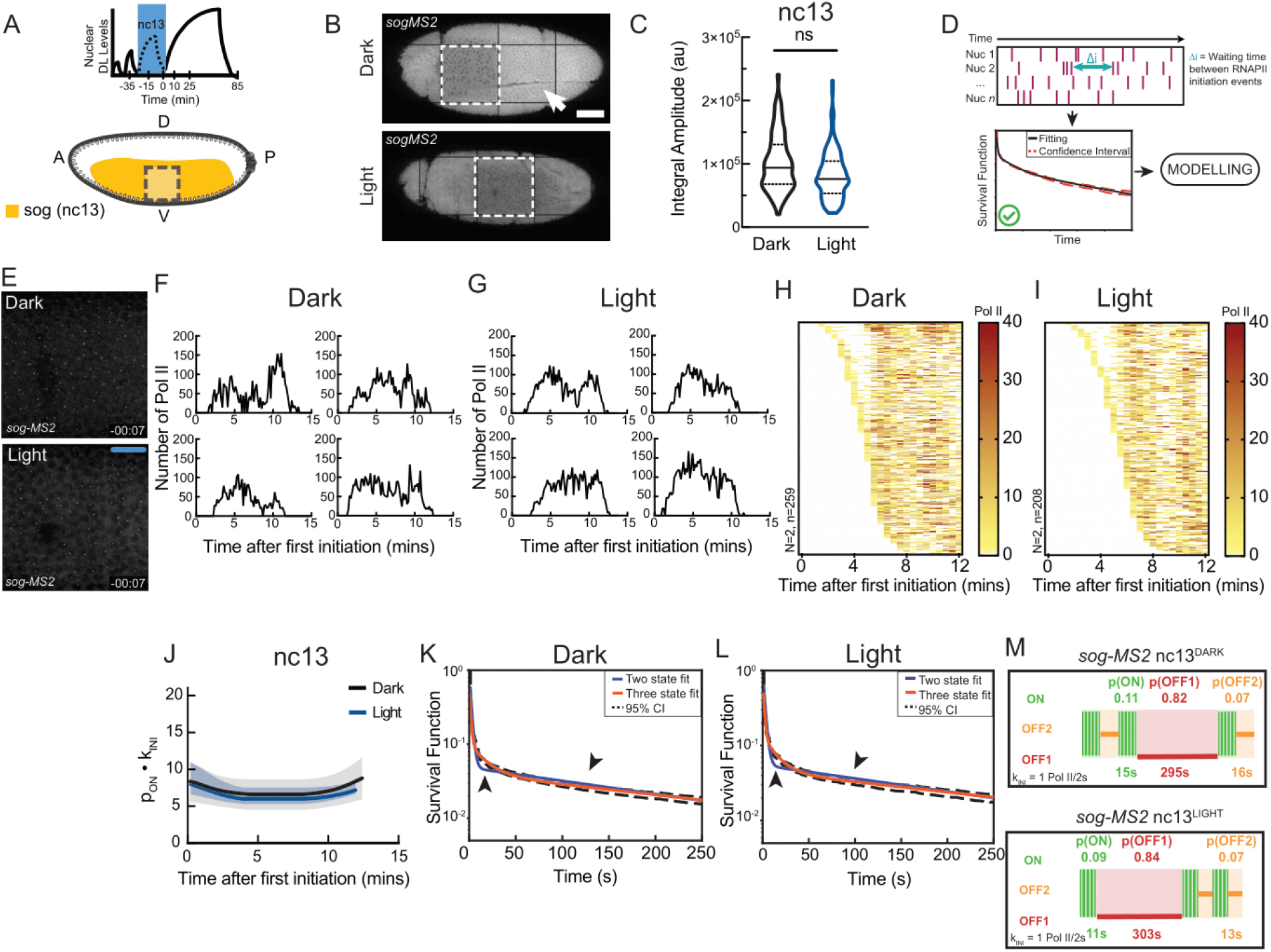
Reduction of DL levels in nc13 does not acutely perturb *sog* bursting kinetics at nc13. **(A)** Schematic of blue light (Dl export) window during nc13. **(B)** Tilescan of late nc14 embryos showing gastrulation in control (upper panel) and attempted gastrulation (lower) after DL export in nc13. Arrowhead indicates invagination associated with gastrulation that is absent following DL export. **(C)** Integral amplitude of *sog-MS2* transcription in nc13 (ns = not significant; Kolmogorov-Smirnov test). **(D)** Schema showing process of model fitting from statistical deconvolution of individual nuclei traces. **(E)** *sog-MS2* transcription in control or DL-exported conditions during nc13. **(F**,**G)** Sample single nuclei *sog* transcriptional traces in nc13 for control (F) and DL-exported (G) conditions. **(H**,**I)** Heatmaps showing the number of polymerase initiation events for *sog* in nc13 in control (H) and DL-exported (I) conditions. Each row represents one nucleus, and the number of Pol II initiation events per 30 s bin is indicated by bin color. **(J)** Kinetic parameter stability as a function of time for *sog-MS2* transcription expressed as the product of the probability to be active (pON) and the RNA polymerase II initiation rate (kINI) in nuclear cycle 13 in control (black) or DL-exported (blue) conditions. **(K**,**L)** Survival function for the distribution of waiting times between polymerase initiation events for *sog-MS2* transcription in control (K) or DL-export (L) conditions. Two-exponential fitting (blue) estimated using the Kaplan–Meyer method extends beyond (arrows) the 95% confidence interval (dashed lines) and is rejected. **(M)** Ideograms of nc13 *sog-MS2* transcriptional kinetics in dark (control) or light (DL-export) conditions. (Live imaging: Dark: N=2 embryos, n=259 nuclei; Light: N=2 embryos, n=208 nuclei).

Since *sog* is expressed in nc13 throughout ventral and lateral regions (the presumptive mesoderm and neurogenic ectoderm) with no respect to the future differentiation of these two presumptive domains, we pooled *sog*^+^ nuclei (with an active TS) for kinetic profiling from the entire imaging window spanning ventral to ventrolateral regions (**Figure 3A,B**, dashed box). Owing to the absence of fluorescently labeled nuclear markers, we were unable to orient live transcription traces relative to mitosis and thus traces were tracked relative to the first detected transcriptional activation event in each nuclear cycle.

As discussed above, the number of active nuclei expressing *sog* increased in nc13 when nuclear DL levels are perturbed with blue light (**Figure 2M**). We also quantified the integral amplitude in nc13 for each trace, which is a proxy for total mRNA output, and noted no difference in total *sog* mRNA output (**Figure 3C**). This was confirmed by quantifying nascent *sog* transcription using single molecule FISH (**Figure S5**), where extended (2h) DL export did not result in substantial perturbations to *sog* nascent transcription compared to the control. Thus, the embryo-level transcription of *sog* appears to be unaffected by optogenetic removal of DL.

However, *sog* transcription is bursty, meaning that the fluctuations, not just the mean mRNA production, may be biologically relevant. Therefore, we next turned to a previously established mathematical modeling approach to examine the underlying kinetic parameters of *sog* transcriptional bursting at the single nucleus level when the DL gradient was perturbed. During a given nuclear cycle, the MS2 signal that carries information on the bursting kinetics, progresses through several stages. Initially, following mitotic exit, no signal is detected and this period corresponds to time for DNA replication and resumption of transcription. This period is typically modeled by considering the distribution of the time to reactivation^28^. Following this initial period, transcription builds up at each transcription foci and can reach a stationary phase, during which the signal remains stable. Bursting during this stationary phase is analyzed using BurstDECONV^29^.

We used BurstDECONV to deconvolve *sog*-*MS2* signal, i.e., reconstruct the sequential polymerase initiation events contributing to this signal (**Figure 3D**)^29^. The deconvolution relies on a model of the contribution of a single polymerase to the MS2 signal, which depends, among other factors, on the length of the transcribed MS2 and post-MS2 sequences. Since the MS2 loops were inserted in *sog’s* first intron, an additional point of consideration for the modeling was to understand the gene’s splicing, recently observed to be largely recursive in human cells^30^. We ensured *sog* splicing was not recursive at this stage by consulting NET-seq data from *Drosophila* embryos^31^. Instead, *sog* splicing likely occurs at intron-exon junctions and is co-transcriptional as supported by FISH data^11^. We therefore considered that the MS2 sequence was entirely transcribed into mature RNA.

Furthermore, the deconvolution method implemented by BurstDECONV assumes that the signal is stationary. Therefore, we first determined whether *sog* transcription in nc13 (**Figure 3E-I**) reached stationary dynamics by plotting the mean waiting time between polymerase initiation events for a short window (81 sec; **Figure 3J**). This represents the mean transcription rate for each window, expressed as the product of the Pol II initiation rate (kINI) in the active state and the probability of a nucleus to be active (PON). Transcription is considered stationary if this value remains stable across sequential time windows. Indeed, this demonstrated that in both dark (nuclear DL, black) and DL-exported (blue) conditions, the underlying kinetic dynamics reached stationarity.

Having established the specific window exhibiting stationarity of the *sog* signal, we next sought to extract *sog* promoter switching rates. To access transcription kinetics, we examined the distribution of waiting times between Pol II initiation events during the first 15 minutes of nc13 and fitted it with a multi-exponential function (**Figure 3K,L**). The multi-exponential fitting gives access to the kinetic parameters driving *sog* transcription such as the duration of the productive (ON) and non-productive (OFF) states as well as the Pol II firing rate in the ON state. Critically, this approach does not presume a specific number of states *a priori*, but instead discovers it as an emergent property of the data itself, resulting from the number of exponentials needed to fit the data. In both dark (nuclear DL) and light (DL export), this distribution could not be fitted by a bi-exponential distribution, indicating that the classical two-state random telegraph promoter model is insufficient to fit the data. A three-exponential fitting, corresponding to a three-state model, was sufficient, however, for both the nuclear and exported DL cases (**Figure 3K,L**, compare blue and orange curves). This three-state promoter scheme consists of a competent ON state and two distinct OFF states, each characterized by a different mean duration: OFF1 (long-lived, 295-303 sec) and OFF2 (short-lived, 13-16 sec). In terms of probability, the prolonged inactive state (OFF1) is the most probable state, dominant in both dark and light conditions. When nuclear DL levels were reduced with blue light in nc13, *sog* promoter bursting dynamics also were captured by a three state model with similar kinetics to those observed in the dark (**Figure 3M**). This analysis of *sog* transcriptional dynamics suggests that upon optogenetic perturbation of DL with the LEXY system in nc13, *sog* transcriptional kinetics are not acutely affected.

### DL export in nc13 leads to mispatterning in nc14, accompanied by a change in bursting kinetics

While in nc13, *sog* is expressed throughout the presumptive mesoderm and neurogenic ectoderm, it is progressively repressed in the nc14 mesoderm via increasing levels of Sna but maintained in the neurogenic ectoderm^32^. Given the specific requirement of DL in nc13 to promote the expression of *sna*, and by extension the repression of *sog* in nc14, we examined the transcription of *sog* in nc14 when nuclear DL levels are perturbed in nc13 (**Figure 4A**).

**Figure 4:**
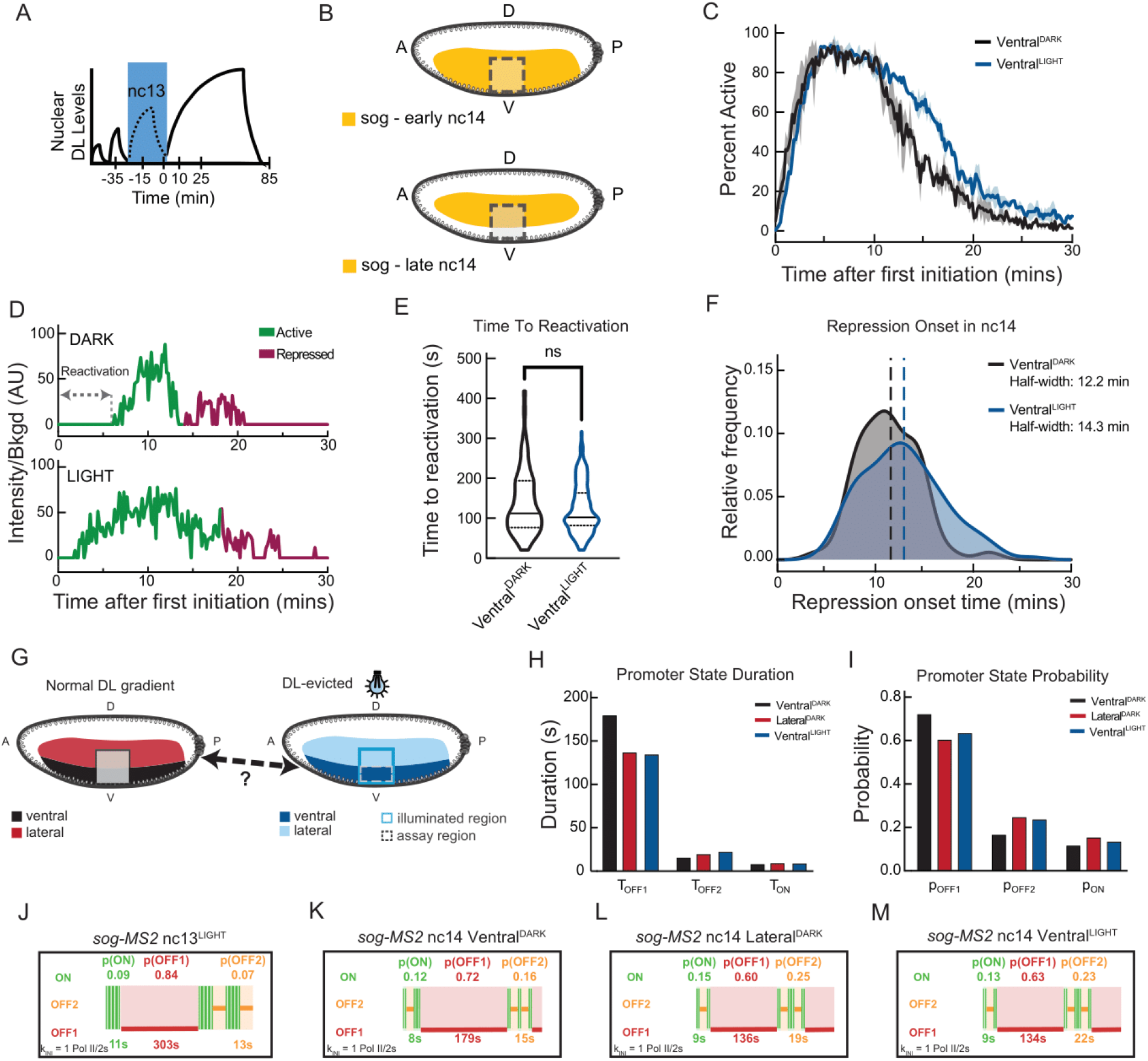
Dorsal export in nc13 is sufficient to transform *sog* kinetics in the mesoderm in nc14. **(A-B)** Schematic of blue light (Dl export) and analysis window (B, dashed lines) for the same movies used in Figure 3 but during nc14 within the indicated embryonic region. **(C)** Fraction of active nuclei in ventral domain in control (black) or DL-exported (blue) conditions, shown as mean ± SEM. Representative insets show relative intensity of *sog* transcription in control (black) or DL-exported (blue) conditions at indicated times. **(D)** Single nuclei traces indicating the time to reactivation and breakpoint between active and repressed paradigms. **(E)** Distribution of time to reactivation for control and DL-exported ventral nuclei (ns = not significant; Kolmogorov-Smirnov test). **(F)** Distribution of breakpoint times for initiation of stable repression in nc14 in control (blue) and DL-exported (red) conditions using Bayesian change point detection. Dashed line indicates median breakpoint time. **(G)** Schema indicating potential equivalency between Lateral^DARK^ and Ventral^LIGHT^ domains. **(H)** Promoter state durations for Ventral^DARK^, Ventral^LIGHT^, and Lateral^DARK^ domains for first 15 minutes of active *sog* transcription in nc14. **(I)** Promoter state probabilities for Ventral^DARK^, Ventral^LIGHT^, and Lateral^DARK^ domains for the first 15 minutes of active *sog* transcription in nc14. **(J-M)** Ideograms of nc13^LIGHT^ and nc14 *sogMS2* transcriptional kinetics in dark (control) or light (DL-exported) conditions. (Live imaging: Dark: N=2 embryos, n=70 nuclei for ventral domain; Light: N=2 embryos, n=162 nuclei for ventral, n=181 nuclei for lateral domain).

Because *sog* is known to have variable expression dynamics based on the tissue in which it is expressed^25,32,33^, we began by examining *sog* repression exclusively in the presumptive mesoderm. In dark embryos, we identified the mesoderm/neuroectoderm boundary based on repression of *sog* by Sna in the mesoderm and spatially selected only those nuclei located in the presumptive mesoderm (see **Figure 4B**). In illuminated embryos, the mesoderm/neuroectoderm boundary driven by Sna protein is perturbed due at least in part to changes in *sna* expression, and thus cannot be used to differentiate the tissues. To circumvent this, embryos were imaged with the presumptive ventral furrow oriented to one edge of the imaging window, and the mesoderm was selected as the 30 µm to either side of the failed furrow at the end of nc14. As expected, given the reduction in *sna* transcription, we observed an extended maintenance of *sog* transcription in the presumptive mesoderm under illumination compared to control dark embryos (**Figure 4C**). *sog* derepression in the ventral part of the embryos was also observed by an alternative approach not relying on MS2/MCP imaging. We performed single molecule RNA FISH in DL^LEXY^ embryos with a more sustained light exposure using an optobox (**Figure S5A-C**). Interestingly, in these conditions the quantification of TS intensities suggests that in the light, transcription of *sog* in ventral nuclei is not different from more laterally located nuclei in the dark (**Figure S5E**).

To understand how reducing DL levels in nc13, and the subsequent reduction of nc14 Sna protein levels (**Figure S1F**), affects *sog* transcription dynamics in nc14, we characterized two metrics from the movies for each nucleus: the time to reactivation (**Figure 4D**, double-arrow domain) and the repression breakpoint (**Figure 4D**, switch from green to red trace), corresponding to the post-mitotic transcription reactivation and to the onset of repression, respectively. We first considered the timing required to reach the first transcriptional activation after mitosis, which we call the time to reactivation^34^ (**Figure 4E**). This time was not significantly different between the control (dark) and DL-exported (light) nuclei. As discussed in Dufourt*, Trullo* *et al*, 2018, the reactivation time has two components, a deterministic component common to all nuclei, and a stochastic component. Although both components contribute to the reactivation period duration, only the stochastic component depends on the number and duration of rate-limiting steps in reassembling the transcriptional machinery^35,36^. The reactivation metric used here accounts only for the stochastic part of *sog* expression in early nc14 when nuclei reinitiate *sog* expression following mitosis. Indeed, time zero is defined here as the moment when the first nucleus resumes transcription after mitosis, rather than the moment of mitosis itself. By making this choice, we exclude the deterministic component of the reactivation time, which is consistent across all nuclei. As mentioned earlier, this approach does not hinder our ability to detect changes in the mechanism of reactivation.

We then turned to Bayesian Change Point Detection (BCPD) to determine the onset of repression, when each individual nucleus underwent a transition from building up repression to stable, full repression^37,38^. For further modeling purposes, the signal is considered stationary after the repression onset. In the DL-exported (light) nuclei, we observed that repression onset for *sog* was significantly delayed (**Figure 4F**, dash lines). Moreover, upon DL-export in 13, *sog* repression within the ventral region appeared much less coordinated (i.e. the repression onset shows greater internuclear variability) in nc14 than in dark conditions (**Figure 4F**). Thus, acute reduction of DL levels in the short temporal window spanning nc13 has a substantial, persistent impact on the dynamics of *sog* transcription later in nc14.

Because DL export in nc13 affects *sna* expression (**Figures 1F and S1F**) and because Sna is an important driver of the mesodermal fate^39,40^, we expect that the DL-depleted ventral cells may have either ‘lost’ or failed to acquire a mesodermal fate and may have adopted instead a more lateral-like fate (**Figure 4G**). To test this hypothesis at the kinetic level, we sought to extract the transcriptional bursting kinetics for *sog* in various spatial regions of the embryo at stationarity in nc14 when kept in the dark or exposed to light at nc13. Ventral region nc14 *sog* data in embryos previously exposed to light in nc13 (Ventral^LIGHT^) or continuously kept in the dark (Ventral^DARK^) were compared with lateral regions (i.e. Lateral^LIGHT^ and Lateral^DARK^). Because embryos fail to gastrulate when exposed to light (i.e. there is limited ventral furrowing), the lateral region was confidently identifiable only in embryos kept in the dark (Lateral^DARK^).

We used BurstDECONV to convert individual fluorescent traces into polymerase initiation events (**Figure S6D-F**) from which kinetic parameters can be modeled^29^ for the three conditions: Ventral^DARK^, Ventral^LIGHT^, and Lateral^DARK^ at stationarity in nc14 (see Methods and **Figure 3D**). After model fitting (**Figure S6J-L**), both the control and DL-exported nuclei could not be fit with a two-state dynamic and instead required a three-state fitting with a competent ON state as well as both long (minutes) and short (seconds) OFF states. In the dark, the *sog* promoter seems to transition between three promoter states in both nc13 and nc14, however the duration and probabilities of the dominant long OFF state are different between these two cycles (**Figures 4H-M and S6M-O**). After DL export in nc13, we observed that the duration of the long non-productive (OFF1) state in nc 14 was reduced by 25% and its probability was decreased by 12% in the ventral domain. Furthermore, the duration of the short non-productive (OFF2) state in nc14 was slightly increased, and its probability was significantly increased by 42% in the ventral domain (**Figure 4H,I**, compare Ventral^DARK^ with Ventral^LIGHT^). Notably, these trends associated with the ventral domain after light exposure closely matched the dynamics of the lateral domain control (**Figure 4H-I**, compare Ventral^LIGHT^ with Lateral^DARK^). Collectively, these results suggest *sog* transcription in the ventral region may convert to a more lateral-like transcriptional profile in nc14 following transient depletion of DL in nc13.

## DISCUSSION

Using an optogenetic approach to control DL localization with LEXY *in vivo* (**Figure 5A**), we identified specific time windows in which DL input is essential for the expression of select DL targets expressed in ventral regions (*sna* and *twi*; **Figures 1** and **5B**). In contrast, we found that lateral genes like *vnd* and *sog* do not respond in the same way as ventral genes to the narrowest critical window of DL activity (i.e. nc13) (**Figures 2** and **5B**). Additionally, we characterized the spatiotemporal bursting kinetics of the endogenous gene *sog* gene across different stages of the patterning process (**Figure 5C**). Notably, when DL is displaced from the nucleus early on, *sog* is derepressed in ventral regions at later stages, resulting in mispatterning. This mispatterning causes ventral cells, the presumptive mesoderm, to instead exhibit properties characteristic of neurogenic ectoderm. These shifts in identity were quantifiable by changes in bursting kinetic properties, with ventral cells adopting a kinetic profile usually associated with lateral cells. These findings were supported by the observation that light exposure at nc13 leads to gastrulation defects, likely because ventral cells can no longer undergo the cell shape changes necessary to support invagination (**Figure 2A,C**).

**Figure 5:**
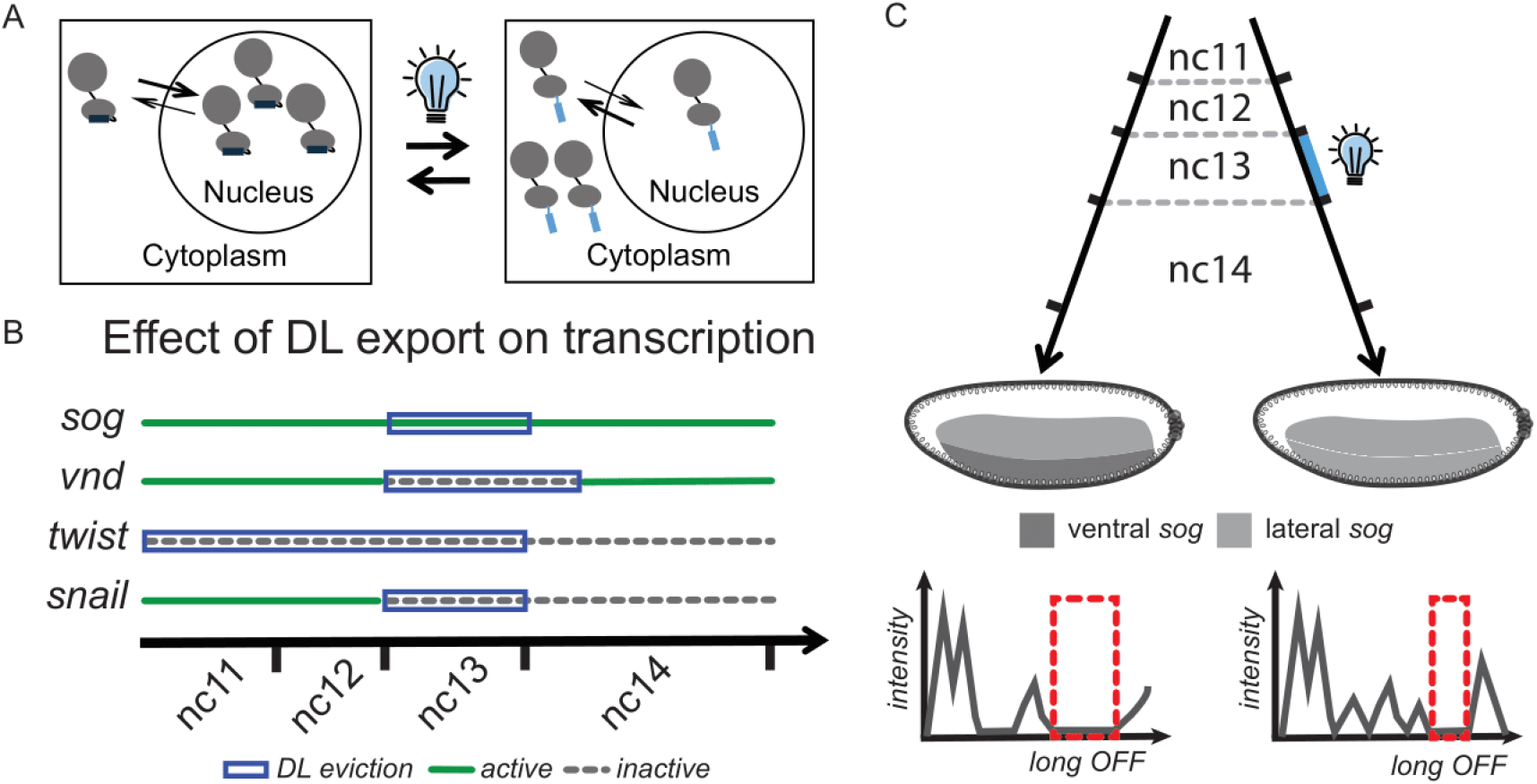
Dorsal export has differential effects on target gene transcription in early embryogenesis. **(A)** Schematic of DL^LEXY^ export dynamics upon blue light stimulation. **(B)** Summary of pattern-level effects of DL export on transcription of known DL target genes when DL is evicted during critical windows (blue box), showing transcription may remain active (green) or be inactive (dashed grey line). **(C)** Summary of single cell-level kinetic switching in ventral *sog* transcription in nc14 after blue light is applied in nc13 (right), demonstrating a change of regime from a mesoderm-like to a lateral-like dynamic (i.e. the long OFF1 state is shorter following light treatment).

These results echo, but are distinct from, the temporal requirement identified for the Bicoid morphogen that controls patterning along the anterior-posterior axis of *Drosophila* embryos. Duration of Bicoid input is important for targets expressed at the anterior pole, presumed to be high threshold targets^4^. However, we found that *sna*, a presumed high-threshold DL target gene, is not sensitive to duration of DL input before mid nc14; as ∼25-30 min blue light exposure during nc12 + nc14 early did not turn off *sna*, whereas a less than 20 min exposure during nc13 did. Instead, our data support the view that *sna* and *twi*, both expressed in ventral regions, require DL input at a critical time (i.e. nc13 for *sna* and nc11-13 for *twi*) when DL activity is necessary so that *sna* and *twi* can be expressed later at nc14. It is notable that *sna* and *twi* exhibit different critical windows of DL input despite being both high-threshold DL targets. We propose this relates to combinatorial control by other TFs in addition to DL, including opposite autoregulatory activities. *twi* is known to exhibit positive autoregulatory feedback whereas *sna* is known to exhibit negative autoregulatory feedback^41^. We suggest that if a little Twi or Sna is made at or following the time that DL is removed at nc13, the respective autoregulatory feedbacks might lead to retention of *twi* and loss of *sna* transcripts. Therefore, the simplistic model of threshold responses to DL is not sufficient to predict whether or not a gene will be retained when DL is perturbed. Previously, we showed that *sna* also exhibits DL dependence at early nc14 using an optogenetic approach based on degradation^18^. Therefore, while DL activity at nc13 is required, it is likely not sufficient for correctly activating *sna*.

There are a number of possible explanations for why DL is critical at nc13. One is that DL may support an epigenetic change or perform some other pioneering activity at the *sna* locus only possible at nc13 that is required for later expression of *sna* at nc14. While it remains unclear why the DL level of mother nuclei at nc13 has an effect in nc14, we can envisage several potential molecular mechanisms including mitotic bookmarking (e.g. GAF)^42^, mitotic-assisted repression^12^, ‘memory’ transcription hubs^43^ or post-transcriptional modulation of mRNA or protein half-life. Our results regarding the criticality of nc13 also agree with a previous study that used an optogenetic approach to activate extracellular signal-regulated kinase (ERK)^44^. Specifically, they found that when light was applied in the trunk at nc13, ectopic expression of *hückebein (hkb)*, a repressor of *sna*, leads to reduction in *sna* transcription and defects in gastrulation.

Our inducible and reversible manipulation of a morphogen TF while recording target gene responses allowed us to dissect time-dependent transcriptional dependencies within a complex gene regulatory network (GRN). Indeed, target gene sensitivity to a dedicated temporal window could be a direct effect or the result of complex feed-forward/combinatorial regulation of other genes by the morphogen input, a common feature of GRNs^45,46^. Gene-gene interactions can influence both the timing and duration of critical temporal windows. Using DL^LEXY^ to decrease DL levels and via our characterizations of critical temporal windows, we can study how the GRN responds to reducing the levels of a specific, lineage-driving TF in a system in which DL levels have recovered to their previous level. For instance, *sna* is a critical component of the gene regulatory network acting to support mesoderm formation and a regulator of neurogenic ectoderm and mesectoderm patterning. Sna protein is known to act as a transcriptional repressor, and without *sna* transcription at nc13 (due to DL export in nc13), Sna protein levels are diminished in nc14, leading to patterning failure. In this manner, we could distinguish both direct, fast effects of DL perturbation (i.e. reduced expression of targets such as *sna, twi*, and *vnd*) as well as phenotypes that present later, after blue light is removed and DL nuclear levels are recovered (i.e. ventral expansion of *vnd* and *sog*).

This study also highlights the value of combining transient perturbations with quantitative approaches to explore more subtle effects on single-cell state changes. We used this insight of the critical time window for *sna* in nc13 to study the transcriptional dynamics of other target genes. Hence, we could decode how DL levels affect bursting across time (nc13 versus nc14) but also across space (ventral versus lateral). Indeed, by nc14 *sog* is repressed in the mesoderm (by Sna) while kept active in lateral regions, thus offering the possibility to compare bursting kinetics in various spatial domains. In both nc13 and nc14 and regardless of DL levels, *sog* promoter dynamics can be recapitulated by a three-state model, comprising a competent ON state, a short-lived OFF state (seconds) and a longer-lived (minutes) highly probable second inactive state (e.g. **Figures 3K,L**, orange line, **and S6J**,**K**,**L**). While our data are unable to demonstrate the biochemical nature of these promoter states, their timescale and their sensitivity to DL levels and spatial region provide clues to discuss what they could represent. The competent ON state, from which polymerase initiates, is generally interpreted as the preinitiation complex (PIC)-assembled promoter state. Importantly, while other studies propose that the probability to be ON stands as the main parameter underlying transcriptional dynamics^8,47,48^, our study uncovered changes in the duration of an inactive state.

The only promoter state that shows a clear change following transient DL manipulation is the longer lived OFF1 state. There are several possible interpretations of this result. In the blastoderm, *sog* expression is orchestrated by two enhancers, an intronic and a distal enhancer, both regulated by DL. Interestingly, deletion studies have suggested that instead of being redundant, these two enhancers integrate activation and repression signals differently^25,33^. For example, the intronic enhancer appears to be the principal enhancer in ventral and ventrolateral regions at nc13. Since the long OFF state changes between nc13 and nc14 (see **Figure 4K-M**), we speculate that this state may correspond to enhancer-promoter interactions: a long OFF state in nc13 when expression primarily relies on one enhancer, that could be reduced in nc14 thanks to the combinatorial action of two enhancers modulated by the presence or absence of a repressor^25^. The kinetics of these long inactive states (minute-range) can possibly provide a clue on the timescale and probabilities of enhancer-promoter encounters^49^. An alternative, possibly simpler explanation, interprets the long OFF1 state as DNA bound repressor molecules. The lifetime of this state in the case of multiple binding sites, representing the time required to clear all the sites, may depend on the number of occupied sites. The concentration of the repressor is thus reflected in this duration, as higher concentrations lead to a longer lifetime.

Taken together our results demonstrate that optogenetic perturbations with high resolution imaging and quantitative modeling can dissect how a TF affects target gene transcription dynamics in space and time. When employed in the context of a pivotal morphogen TF, patterning can be disrupted and lead to mispatterning: here, presumptive mesoderm (ventral) to neurogenic ectoderm (lateral), detectable at the level of the kinetics of transcriptional bursting.

## MATERIALS & METHODS

### Drosophila stocks

For determining the critical window of DL action, crosses were kept at 18°C. *dl-LEXY; nos>MCP-mCherry-nls/TM3* virgins were crossed to *twi-MS2, sna-MS2, vndEEE-MS2/CyO*, or *sog-MS2/y* males (see Table S1). MS2 was added to the endogenous locus to generate *twi-MS2* (this study) and *sog-MS2*^*1*^ using CRISPR/Cas9. *sna-MS2* is a large (∼25 kb) reporter construct, which was inserted on the third chromosome, and contains the known regulatory sequences of *sna*, MS2 at the 5’UTR, and part of the yellow gene^23,51^. *vndEEE-MS2* is a reporter line containing the *vndEEE* enhancer, the *eve* promoter, MS2 at the 5’UTR, and lacZ^24^. *dl-mCh-LEXY* mothers were used to image DL^mCh-LEXY^ levels.

In all other cases, stocks and crosses were maintained at 25°C. *dl-LEXY; nos>MCP-RFPT* stocks were maintained in the dark as a double homozygous line. Two types of *sog*-*MS2* strains were used, both CRISPR alleles containing 24MS2 stem loops in the first intron. Experiments described in Figure 2 used a *sog*-*MS2* stock described in McGehee & Stathopoulos^1^ and *dl-LEXY; nos>MCP-mCherry-nls/TM3*; while data from Figure 3 and 4 used a *sog*-*MS2* allele described previously^25^ and *dl-LEXY; nos>MCP-RFPT*. For live imaging, *dl-LEXY; MCP-mCherry/TM3* (or *MCP-RFPT*) females were crossed to *sog*-*MS2* males. For single molecule FISH, *dl-LEXY/dl-LEXY* homozygous flies were used for embryo collections.

### Live Imaging and Quantification to Determine Critical Windows

Live imaging for quantification of nascent transcription spots (MCP/MS2 puncta) or DL^mCh-LEXY^ levels was performed as described previously^1^. Briefly, embryos were collected at 18°C and then hand dechorionated in the dark, using a red film (Neewer, 10087407). Embryos were oriented and then transferred to a coverslip containing heptane glue. Embryos were covered with water to prevent desiccation. Imaging occurred on a Zeiss LMS 800 using a 25x immersible objective (LCI Plan-Neofluar 25x/0.8 Imm Korr DIC M27) at 1.7 zoom and at 20-21°C. A 561 nm laser at 1% laser power with 800 V gain on a GaAsP PMT detector was used to detect the MCP-mCherry signal. A 488 nm laser at 4.5% laser power was used to perform blue light illumination. Images consist of 30 z-planes per timepoint at 1 um thickness. Images were taken every ∼25 seconds, starting as soon as the previous z-stack finished. Images were captured as 16 bit, with 512 by 512 pixel resolution, with each pixel being 0.29 um in length and width. In addition to the previous settings, eight *sna-MS2* movies were taken using slower settings. Images were taken every ∼2 min and each z-slice was 1024 by 1024 pixels with each pixel being 0.15 um long. Imaging was terminated after observing gastrulation or movement of the nuclei congruent with germ band extension when gastrulation was absent or not observable. Spots were detected as described previously^1^. Briefly, the image is filtered using a median filter, the background is subtracted, and the image is blurred using a Gaussian filter. Then two thresholds are applied to detect the active sites of transcription and remove noise. The embryo is segmented using a different threshold and any spots detected outside the embryo are removed. This is performed for individual time frames, such that spots are counted instantaneously and not cumulatively. This allows for visualizing decreases in the number of active sites of transcription.

The same imaging scheme was used to image the DL^mCh-LEXY^, except the 561 nm laser was used at 2% laser power. To quantify the DL^mCh-LEXY^ levels we used a similar method as described previously^1^. Briefly, average intensity projections were segmented using MATLAB’s edge function and a Laplacian of Gaussian with a standard deviation of four for the filter. Water-shedding was performed using MATLAB’s watershed function to disconnect any nuclei that were connected. Objects were filtered by size to remove large (i.e. two connected nuclei) or small (i.e. non nuclei or partial nuclei) objects. Additionally detected objects were filtered by intensity. Specifically, images were background subtracted and an intensity threshold was applied. Objects that overlapped in both the intensity thresholded mask and the Laplacian of Gaussian mask were kept. The embryo was detected by blurring the image, and using a normalized threshold of 0.005, followed by morphological opening and closing. The boundaries of the embryo were detected and used to fit an ellipse. From the ellipse, the midline was determined and only nuclei within 100 pixels above or below the midline were included for analysis. The average intensity was calculated for each individual nucleus at a given time point and then was averaged together for each time point and plotted as the mean ± s.d.

### Live Imaging to Study Bursting Kinetics

Embryos were permitted to lay for 2 h in the dark at 25°C prior to collection for live imaging. All preparation was performed in the dark with minimal illumination through a red filter. Embryos were hand dechorionated and mounted on a hydrophobic membrane prior to immersion in oil to prevent desiccation and addition of a coverslip. Live imaging was performed with an LSM 880 with Airyscan module (Zeiss). Z-stacks comprised of 30 planes with a spacing of 0.5 μm were acquired at a time resolution of 10.21 s per stack in fast Airyscan mode with laser power measured and maintained at 14.5 uW using a ThorLabs PM100 optical power meter (ThorLabs Inc.). All movies were performed with the following settings: constant LEXY export by a 488-nm laser and RFP excitation by a 561 nm were captured on a GaAsP-PMT array with an Airyscan detector using a 40x Plan Apo oil lens (NA = 1.3) at 2.0 zoom on the lateral region of the embryo from the presumptive mesoderm to the dorsal border of the *sog* domain. Resolution was 104.7 um x 104.7 um with bidirectional scanning. Airyscan processing was performed using 3D Zen Black v3.2 (Zeiss).

### Live Imaging Analysis for Transcriptional Kinetics

Data was acquired in TZXY for each channel. A custom software was developed in Python™ to detect and track MS2/MCP puncta in the absence of a nuclear marker.

Briefly, Airyscan-processed data is loaded, concatenated and maximum intensity projected for visualization purposes. To detect transcription puncta, raw data was filtered with a 3D Laplacian of Gaussian Filter followed by manual thresholding based on the mean and standard deviation of the filtered image. Rare cases of optically-resolvable sister chromatid separation was resolved by merging objects with a distance smaller than a user-defined threshold (generally 1-2 px). A volume threshold was applied to remove false detection events due to noise. Finally, spots were tracked through time using a 2D minimal distance criteria with a threshold distance to block erroneous associations. A gap of 8 frames or less during tracking with a spot reappearing within the distance threshold was considered a continuous trace, while gaps of more than 8 frames were considered new detection events. Traces that were artificially short relative to the length of the nuclear cycle were eliminated.

Following tracking, background estimation was performed in 3D for each independent frame as the average intensity value of the pixels surrounding the spot. Final spot intensities were then expressed as their values divided by this estimated background to ameliorate technical error such as laser power fluctuation and z-dependent fluorescence intensity changes. This rescaling technique is sufficient unless photobleaching is too high. To monitor overall photobleaching, the overall behavior of the intensity on all the 4D stack excluding the transcriptional puncta was estimated, as their intensity variation is driven by biological factors. All non-signal pixels time frame per time frame were summed and used to estimate bleaching, which was measured to reach 30% at the end of 30 minutes (**Figure S1F**). In nc13, the entire imaged region was used for analysis, and in nc14 the imaged region was divided into either ventral or lateral domains based on the presence of the mesoderm/neurogenic ectoderm border (dark control) or the position of the failing gastrulation furrow (DL-exported).

### Bleaching analysis

To monitor photobleaching, the overall behavior of the intensity on all the 4D stack excluding the transcriptional puncta was estimated, as their intensity variation is driven by biological factors. The intensity of all non-signal pixels throughout nuclear cycle 14 were summed independently for each time point and movie, and fitted with an exponential decay function. For spot counting (*twi-MS2*, **Figure S3A**) and *dl-mCh-LEXY* (**Figure S2E**) movies, bleaching was analyzed for 30 minutes following the peak of the background signal in nc14. For *sog-MS2* bursting movies (**Figure S3B**), bleaching was analyzed from the onset of nc14.

### Single molecule Fluorescence In Situ Hybridization (smFISH)

*dl-LEXY* homozygous flies were permitted to lay for 2 hours in the dark, followed by incubation of the embryos in blue light (30% power, 0.67 duty cycle) for a further two hours in an optobox, or incubation for a further two hours in the dark as appropriate. Embryos were fixed in 10% formaldehyde/heptane under blue light or in the dark as appropriate for 25 min with shaking before a methanol quench and stored at −20°C in methanol before use.

smFISH probes targeting *twist* conjugated to Quasar-580 (LGC Biosearch Technologies) were designed and used as previously described^28^. smiFISH probes targeting the 5’ region of the *sog* transcript were designed using Oligostan (Integrated DNA Technologies, Inc.)^52^. Probes were resuspended in TE at equimolar concentrations. Embryos were prepared for smiFISH as previously described^53^ before mounting in ProLong Gold mounting media (Life Technologies).

### Immunofluorescence

*dl-LEXY/dl-LEXY* homozygous embryos were collected as described above for smFISH. Embryos were progressively rehydrated in PBT before blocking in PBT and 0.5% BSA, followed by incubation overnight at 4°C with mouse anti-Dorsal (1:500; Developmental Studies Hybridoma Bank) and rabbit anti-Snail (1:800; this study). Embryos were washed four times in PBT followed by two hours of incubation with donkey anti-mouse AF-488 (1:1000; Life Technologies), donkey anti-rabbit AF647, and DAPI (1:1000). Embryos were washed four times in PBT before mounting in ProLong Gold mounting media (Life Technologies).

### Fixed Sample Imaging and Analysis

For smFISH, imaging was performed on an LSM 880 with Airyscan module (Zeiss). Z-planes were acquired with 0.3μm spacing spanning the entire nuclear volume, using laser scanning confocal in Airyscan super-resolution mode with a zoom of 3.0. DAPI excitation was performed with a 405-nm laser, AF-488 excitation with a 488-nm laser, and Q580 excitation with a 561-nm laser. Detection was performed using a GaAsP-PMT array coupled to an Airyscan detector. Laser power was maintained between all channels by intensity measurement prior to imaging (ThorLabs, Inc.). Airyscan processing was performed using 3D Zen Black v3.2 (Zeiss) prior to analysis. Embryos were staged based on membrane invagination.

Analysis was performed using previously published custom software^28^. For DL nuclear level analysis (**Figure S1D**), a post-processing tool was developed to spatially define rows of nuclei orthogonal to the dorso-ventral axis by isolating the centroid of each nucleus and defining the number of bins based on the square root of the total number of nuclei in the image. Nuclei were clustered into bins based on the appropriate dorso-ventral axis coordinate of the centroid. For smFISH analysis, mesoderm nuclei were defined as having *twi*^*+*^ transcription site signal in the nucleus.

### Mathematical modeling

To ensure a stable transcriptional dynamic and to avoid any confounding factors due to bleaching during nc14, we limited the analysis window to the first 15 minutes after active transcription began (**Figure S4G-I**). We estimate this corresponds to the first 20-22 minutes after mitosis based on the post-mitotic reactivation window from previously published live imaging studies of *sog* in nc14^25^. For nc13, the entire nuclear cycle was retained for analysis as transcription reached and maintained a steady state (Figure 3J).

Subsequently, the analysis was conducted in three parts. The first part involved determining the distribution of postmitotic reactivation times. The reactivation time has two components: a deterministic component, which is the same for all nuclei, and a stochastic component, which varies among nuclei. Like in Dufourt et al. 2018^34^, we have estimated the stochastic component by setting as time origin the time when the first nucleus activates.

The second part involved determining repression onset time detection. For the determination of this time we have used a Bayesian Change Point Detection (BCPD) method. This method, introduced in Adams and Mackay 2007 models the statistics of the signal and identifies changepoints, i.e., moments when the signal’s statistical properties change^38^. The BCPD method has already been tested on time-inhomogeneous live transcription in *Drosophila* embryos^37^.

The third part uses BurstDECONV^29,54^ to determine the transcriptional bursting kinetic parameters for stationary segments of the signal. When the signal is repressed, the signal segment after repression onset is considered stationary. BurstDECONV first deconvolves the MS2 signal and infers the positions of the transcription initiation events. The distribution of waiting times between successive events is modeled as a multi-exponential distribution. Finally, the parameters of the multiexponential distribution are used to calculate the kinetic parameters of Markovian transcriptional bursting models assumed to function at stationarity. This method has been tested on live transcription data from *Drosophila* embryos and human cells as well as on synthetic datasets^29,53,54^ and has rigorous probabilistic and algebraic foundations^55^.

For the extraction of kinetic parameters, it was imperative to ensure that we were in a steady-state regime. To study the homogeneity of the signal, we estimated the mean waiting time between successive polymerase initiation denoted <τ> across a narrow moving window. This value, for each window, is related to the product of pON, the probability that the promoter is ON, and kINI, the initiation rate in the ON state. Therefore, if <τ> is constant across different time windows, then the product of pON (the probability that transcription is active) and kINI (the transcription initiation rate in the active state; here it is assumed that initiation does not occur in the inactive states) is constant, suggesting that the kinetic parameters are constant (see Pimmett et al., 2024 for more details)^37^. The width of the moving window used here is 8 frames, or 82 sec. We used this moving window to show that the kinetic parameters are constant on a maximal duration of 15 minutes after the first activation of the first nuclei in each movie. Finally, using time series of this maximal duration, we were able to apply our existing pipeline to extract kinetic parameters and perform transcriptional bursting model selection^29^. The hyperparameters used for the BurstDECONV pipeline are: a RNA Polymerase II speed of 25 bp/sec (equivalent to a dwell time of 414 sec), a time resolution of 10.21, and distances between the transcription start site and the start of MS2 loops, the size of the MS2 loops, and the distance between the end of the MS2 loops and the 3’ splice site of 2553 bp, 1267 bp, and 9081 bp, respectively. Given the insertion of the MS2 sequence into the intron, persistence of the MS2 signal at the TS will be reduced if the intron is removed via splicing, thus impacting the signal deconvolution ^56^. We ensured *sog* splicing was not recursive at this stage by consulting NET-seq data from *Drosophila* embryos ^31^. Instead, *sog* splicing likely occurs at intron-exon junctions and co-transcriptionally as supported by FISH data^11^. We considered both two-state (random telegraph) and three-state Markovian transcriptional bursting models as shown in **Figures 3K,L and S4J**,**K**,**L**.

## ACKNOWLEDGEMENTS

We are grateful to Eileen Furlong and Alessandro Dulja (EMBL, Heidelberg) for help with setting up the optobox, Sarah Bray, Hernan Garcia, and Chris Rushlow for providing fly stocks, Mike Levine for providing plasmid DNA, and Louise Maillard and Leslie Dunipace for comments on the manuscript. We also thank Pedro Prudêncio (Carmo Fonseca lab) and Amal Makrini (IGMM, CNRS) for the analysis of published NET-seq data. We acknowledge the Montpellier Ressources Imagerie facility (France-BioImaging), the Biocampus *Drosophila* facility of Montpellier, and the Caltech Beckman Institute Imaging Facility. This study was supported by funding from the National Institute of Health grant R35GM118146 to A.S as well as ERC SyncDev and ANR HubDyn to ML. M.D is supported by the CNRS and University of Chicago Joint PhD programme. V.P was initially supported by ERC syncDev and then ANR HubDyn. M.L., O.R., and A.T. are sponsored by CNRS.

## AUTHOR CONTRIBUTIONS

A.S., M.L, V.P., and J.M. conceived the project and planned the experimental approach. A.S. and M.L. directed the project. J.M. and V.P. performed all experiments, and J.M, VP, and A.T. performed all quantitative analysis of imaging data. A.T implemented the OptoTrack code, suitable for image analysis in the absence of a nuclear marker. M.D and O.R performed mathematical modeling. All authors analyzed and interpreted the data. The manuscript was written by V.P., J.M., A.S., and M.L with edits provided by O.R, M.D and A.T.

## SUPPLEMENTARY INFORMATION

### Supplemental Figure Legends

**Supplemental Figure 1: Use of LEXY system for optogenetic regulation of DL nuclear localization**. Related to Figures 1 and 3. **(A)** Schema of the LEXY system ^19^ in the dark (top) and during blue light exposure (bottom). **(B)** Diagram of the *dl-LEXY* and *dl-mCh-LEXY* loci showing insertion of the LEXY and mCh-LEXY tags (blue) into the DL (yellow) C-terminus ^1^. **(C)** Schema of dark (left) and light-exposed (right) embryos demonstrating a reduction in ventral nuclei of DL^LEXY^ in the presence of light (“+hν”) versus its absence (“-hν”). **(D)** Representative sagittal images of dorsal and ventral nuclei in control and light-exposed embryos showing DL nuclear levels (green) in nc14 after 2h of light exposure. Nuclear DL signal was quantified relative to ventral dark control. Under blue light, likely some low DL levels remain in the nucleus. **(E)** Representative sagittal Z-planes in dark control and light-exposed embryos immunostained with anti-DL and anti-Sna antibodies demonstrating nuclear DL (green) and Sna (purple) signals are reduced upon illumination. **(F)** Representative maximum intensity projections in dark control and light-exposed embryos immunostained with anti-Sna antibody (green) and DAPI (blue) demonstrating loss of Sna protein in mid-nc14 upon illumination. Scale bar is 100um.

**Supplemental Figure 2: Dorsal levels recover in nc14 when removed at nc13 with blue light**. Related to Figure 1. **(A-B)** DL^mCh-LEXY^ signal in the dark (A) or with blue light at nc13 (B) in embryos laid by mothers homozygous for *dl-mCh-LEXY*. **(C)** Nuclei were segmented using the DL^mCh-LEXY^. When no signal was present (during division or under blue light) the intensity was set to zero. The average intensity for nuclei were then averaged for each individual embryo and plotted. **(D)** The same as in (C) except levels were normalized by dividing by the maximum average intensity at nc12. nc12 was chosen so that embryos were normalized to the DL^mCh-LEXY^ signal intensity before blue light exposure. The black lines are in the dark and the blue lines are with blue light at nc13 (mean ± s.d.). Comparing the normalized levels in nc14 between dark and light at nc13, there is little difference, suggesting that DL levels recover at nc14 when blue light is used to remove Dorsal from the nucleus at nc13. **(E)** Quantification of DL^mCh-LEXY^ bleaching in living nc14 embryos for dark (black) and light-exposed (blue) embryos. N=3 embryos for all conditions.

**Supplemental Figure 3: Bleaching quantification during nc14 does not strongly impact signal detection**. Related to figures 1 and 3. **(A)** Quantification of MCP-mCherry-NLS bleaching in living nc14 embryos for dark (black) and light-exposed (blue) embryos. **(B)** Quantification of MCP-RFPT bleaching in living nc14 embryos for dark (black) and light-exposed (blue) embryos. N=2 embryos for all conditions.

**Supplemental Figure 4: *vnd* levels recover in nc14 when illuminated at nc14 for 10 min**. Related to figure 2. **(A)** The *vndEEE-MS2* reporter construct used for tracking *vnd* expression. **(B)** Schematic of the field of view used to image ventrolaterally. **(C**,**D)** *vndEEE-MS2* at nc14 in the dark (C) or with 10 min of blue light (D). **(E)** Quantification of the number of active TS detected for *vndEEE-MS2*. The black line is in the dark and the blue line is with blue light at nc14 (mean ± SEM; n=3 embryos for each). Foci are circled in orange for *vnd*. t=0 indicates the start of imaging in mid nc14.

**Supplemental Figure 5: Quantification of the effects of DL nuclear export on dorsoventral patterning in fixed embryos**. Related to Figures 2 and 3. **(A)** Schema of custom-built optogenetic manipulation system^20^. **(B)** Illumination schema for control and DL-exported embryos. **(C)** Representative smiFISH maximum intensity projections of the nuclear volume for nc13, nc14a and nc14c *dl-LEXY/dl-LEXY* embryos in control (dark) and DL-exported (light) conditions. Mesoderm is indicated by a dashed box. **(D)** Quantification of *sog* TS intensity for nc13 embryos in control and DL-exported conditions. **(E)** Quantification of *sog* TS intensity for the indicated domains and light conditions from a representative randomly selected sample of 200 nuclei for each domain/condition. **(F-G)** Quantification of *sog* TS intensity in early nc14 for ventral (F) and ventrolateral (G) domains. Significance determined by Kruskal-Wallis test.

**Supplemental Figure 6: *sog* bursting kinetics in nc14 after DL export in nc13**. Related to Figure 4. **(A-C)** Sample single nuclei *sog* transcriptional traces during the first 30 minutes of nc14 for Ventral^DARK^ (A), Ventral^LIGHT^ (B) and Lateral^DARK^ (C) regions. Trace colors indicate active (green) and repressed (purple) phases as determined by Bayesian Change Point Detection. **(D-F)** Heatmaps showing the number of polymerase initiation events for *sog* in nc14 for Ventral^DARK^ (D), Ventral^LIGHT^ (E) and Lateral^DARK^ (F) regions as a function of time. Each row represents one nucleus, and the number of Pol II initiation events per 30 s bin is indicated by bin color. **(G-I)** Kinetic parameter stability as a function of time for *sog-MS2* transcription, expressed as the product of the probability to be active (pON) and the RNA polymerase II initiation rate (kINI) in nuclear cycle 14 in nc14 for Ventral^DARK^ (G), Ventral^LIGHT^ (H) and Lateral^DARK^ (I) regions. **(J-L**) Survival function for the distribution of waiting times between polymerase initiation events for *sog* transcription in nc14 for Ventral^DARK^ (J), Ventral^LIGHT^ (K) and Lateral^DARK^ (L) regions. Two-exponential fitting (blue) estimated using the Kaplan–Meyer method extends beyond the 95% confidence interval (dashed lines) and is rejected. **(M-O)** Survival function fitting indicating most parsimonious kinetic parameters and boundaries of the 95% confidence interval for the indicated domains.

### Movie Legends

**Movie S1**. *twi-MS2* (magenta) in *dl-LEXY* in the dark, with light at nc11-13, and with light at nc13. Related to Figure 1. A ventral view is shown in all panels. Nascent transcription was only detected above a threshold for Movies S1-S3, and spots can only reliably be discerned before the embryo initiates gastrulation.

**Movie S2**. *sna-MS2* (magenta) in *dl-LEXY* in the dark, with light at nc13, and with light at nc12 & 14. Related to Figure 1. A ventral view is shown in all panels.

**Movie S3**. *vndEEE-MS2* (orange) and *sog-MS2* (yellow) in *dl-LEXY* in the dark and with light at nc13. Related to Figure 2. A ventral view is shown in all panels.

**Movie S4**. *sog-MS2* in *dl-LEXY* in the dark and with light at nc13, related to Figures 3 and 4. Views relate to ventrolateral regions of embryos as shown in Figure 3B.

**Table S1:** Reagents and software used in this manuscript.

| REAGENT or RESOURCE | SOURCE | IDENTIFIER |
| --- | --- | --- |
| **Antibodies** |  |  |
| Mouse Anti-Dorsal | Developmental Studies Hybridoma Bank (DSHB) <sup>57</sup> | Cat# anti-dorsal 7A4; RRID: AB_528204 |
| Rabbit anti-Snail | This study | N/A |
| Alexa Fluor 488 donkey anti-mouse | Thermo Fisher Scientific | Cat# A21202; RRID: AB_141607 |
| Alexa Fluor 647 donkey anti-rabbit | Thermo Fisher Scientific | Cat# A31573; RRID: AB_2536183 |
| **Experimental Models: Organisms/Strains** |  |  |
| *D. melanogaster*. w; dl-LEXY/CyO; PrDr/TM3 | McGehee & Stathopoulos <sup>1</sup> | N/A |
| *D. melanogaster*. w; dl-mCherry-LEXY/CyO | McGehee & Stathopoulos <sup>1</sup> | N/A |
| *D. melanogaster*. w; dl-LEXY/CyO; MCP-mCherry (w+, NLS)/TM3 | McGehee & Stathopoulos <sup>1</sup> | N/A |
| *D. melanogaster*. w; dl-LEXY; MCP-RFP<sub>t</sub> | This study | N/A |
| *D. melanogaster*. sna-MS2 BAC (III) | Irizarry et al <sup>18</sup> | N/A |
| *D. melanogaster*. sog-MS2 (I); Sp/CyO | McGehee & Stathopoulos <sup>1</sup> | N/A |
| *D. melanogaster*: *sog*-MS2 (I) | Whitney et al <sup>25</sup> | N/A |
| *D. melanogaster*: *w*; *twi*-MS2 (II) | This study | N/A |
| *D. melanogaster*: *w*; *P*{*w*[+*mC*]=*vnd*EEE-*peve*-24xMS2-*lacZ*-SV40}*attP*40 | Falo-Sanjuan & Bray <sup>24</sup> | N/A |
| *D. melanogaster*: *y*[1] *M*{*w*[+*mC*]=*nanos*-*Cas9*.*P*}*ZH*-2A *w*[*] |  | RRID:BDSC_54591 |
| **Oligonucleotides** |  |  |
| *sog* smiFISH probes | This study | N/A |
| FlapY_AF488 | Integrated DNA Technologies | N/A |
| *twi* smFISH probes | LGC Biosearch Technologies | N/A |
| **Recombinant DNA** |  |  |
| pHD-DsRed | Gratz et al <sup>58</sup> | Addgene 51434 |
| Plasmid: *twi*-MS2, DsRed homologous repair | This study | N/A |
| **Software and Algorithms** |  |  |
| Zen 3.0 (Blue edition) | Zeiss | N/A |
| Zen 3.2 (Black edition) | Zeiss | N/A |
| Fiji/ImageJ | Schindelin et al <sup>59</sup> | https://imagej.nih.gov/ij/ |
| Prism 10 | Graphpad | https://www.graphpad.com/ |
| McGehee_2024 | McGehee & Stathopoulos <sup>1</sup> | https://github.com/StathopoulosLab/McGehee_2024 |
| run_analyze_xs | Trisnadi et al. <sup>60</sup> | https://www.sciencedirect.com/science/article/pii/S1046202312002629?via%3Dihub |
| OptoTRACK | This study | https://github.com/ant-trullo/OptoTrack |
| BCPD analysis | Pimmett et al <sup>37</sup>; Adams & MacKay <sup>38</sup> | https://github.com/mariadouaihy/BCPD_inhomogeneous_transcriptional_signal |
| BurstDECONV | Douaihy et al <sup>29</sup> | https://github.com/oradules/BurstDECONV |
| Oligostan | Tsanov et al <sup>52</sup> | https://bitbucket.org/muellerflorian/fishquant |
| smFisher | Dufourt et al <sup>28</sup> | https://github.com/ant-trullo/smFiSH_software<br>https://github.com/ant-trullo/StripesAnalysisStudy |

