## Supplemental Figures 1-6 for "Optogenetic manipulation of nuclear Dorsal reveals temporal requirements and consequences for transcription"

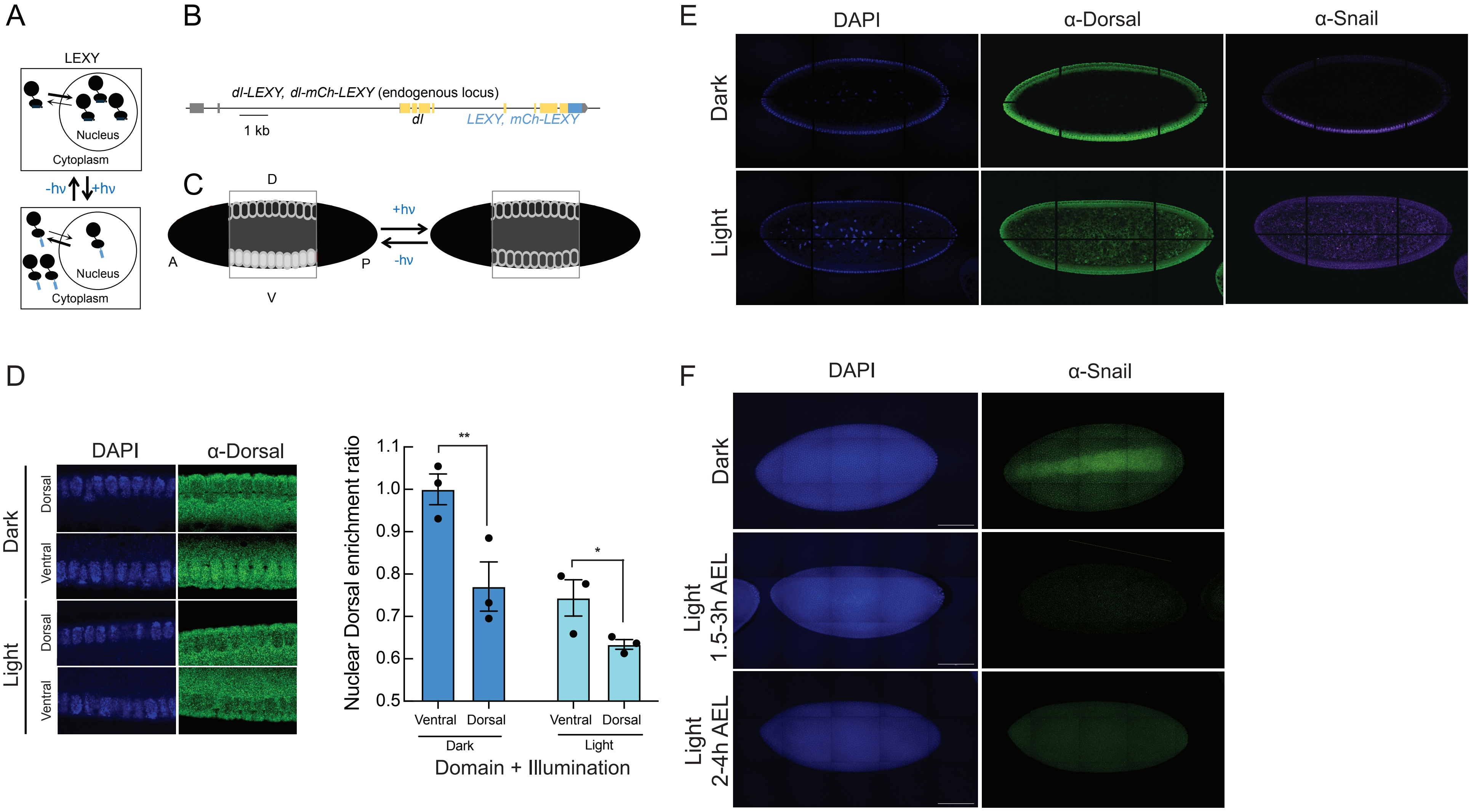
A
LEXY
Nucleus
Cytoplasm
-hν
+hν
Nucleus
Cytoplasm
B
dl-LEXY, dl-mCh-LEXY (endogenous locus)
1 kb
dl
LEXY, mCh-LEXY
C
D
A
P
V
+hν
-hν
E
DAPI
α-Dorsal
α-Snail
Dark
Light
D
DAPI
α-Dorsal
Dark
Light
Dorsal
Ventral
Dorsal
Ventral
Nuclear Dorsal enrichment ratio
1.1
1.0
0.9
0.8
0.7
0.6
0.5
**
*
Ventral
Dorsal
Ventral
Dorsal
Dark
Light
Domain + Illumination
F
DAPI
α-Snail
Dark
Light 1.5-3h AEL
Light 2-4h AEL

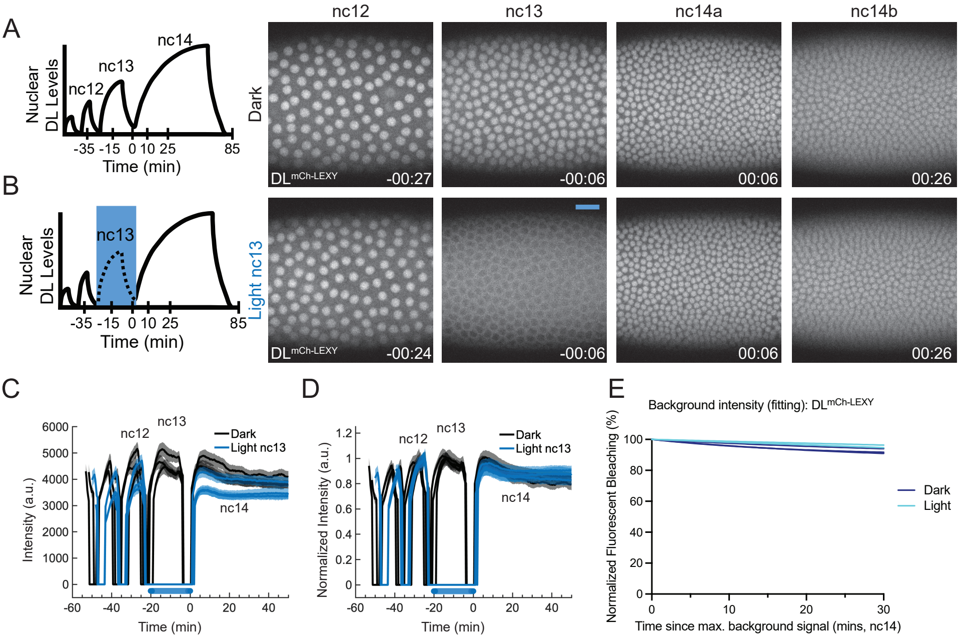

A
B
C
D
E
nc12
nc13
nc14a
nc14b
Nuclear DL Levels
Time (min)
-35 -15 0 10 25 85
nc14
Dark
Light nc13
DL mCh-LEXY
-00:27
-00:06
00:06
00:26
-00:24
Intensity (a.u.)
Normalized Intensity (a.u.)
Light nc13
Background intensity (fitting): DL mCh-LEXY
Normalized Fluorescent Bleaching (%)
Time since max. background signal (mins, nc14)
Light

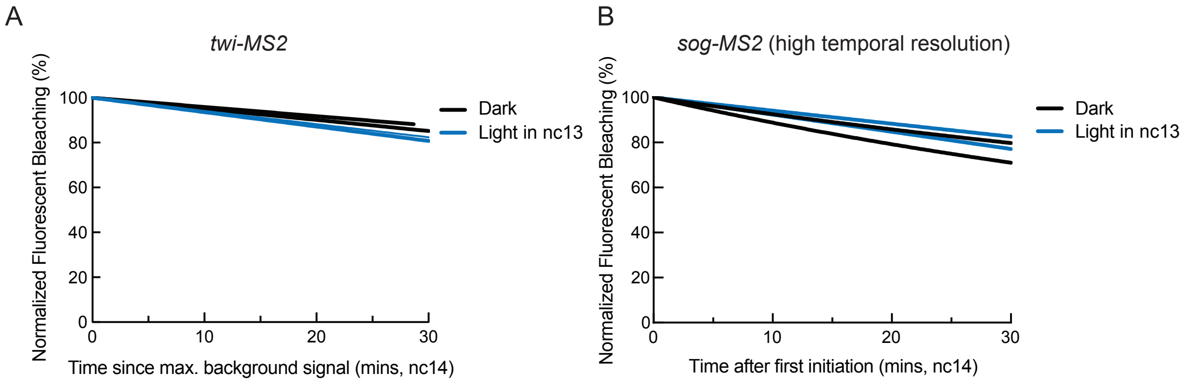
A
twi-MS2
Normalized Fluorescent Bleaching (%)
100
80
60
40
20
0
0
10
20
30
Time since max. background signal (mins, nc14)
Dark
Light in nc13
B
sog-MS2 (high temporal resolution)
Normalized Fluorescent Bleaching (%)
100
80
60
40
20
0
0
10
20
30
Time after first initiation (mins, nc14)
Dark
Light in nc13

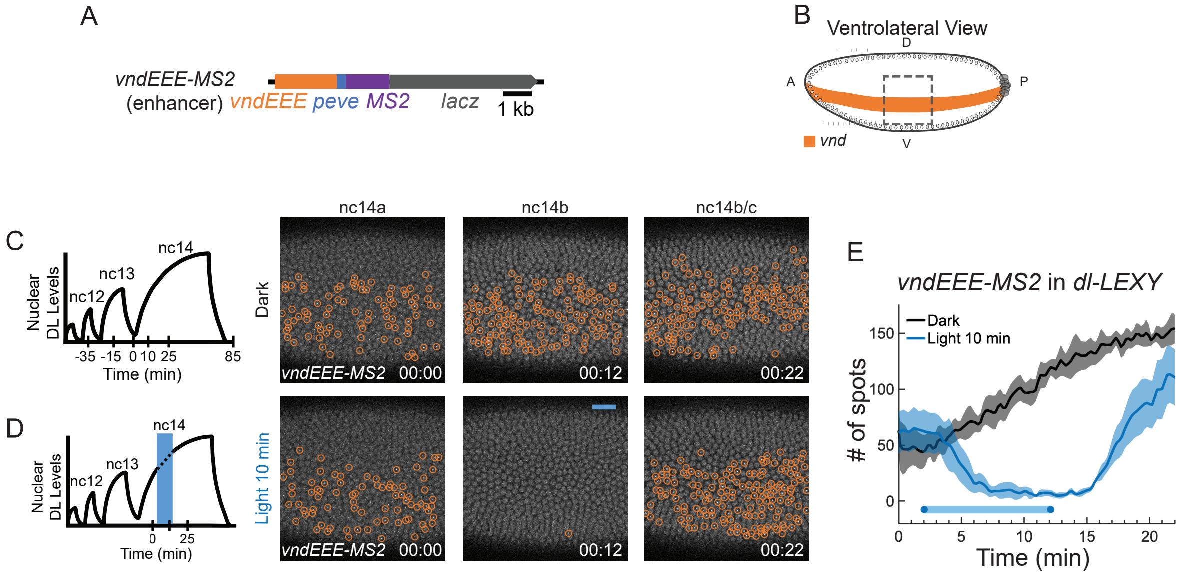
A
vndEEE-MS2
(enhancer) vndEEE peve MS2 lacz
1 kb
B
Ventrolateral View
D
A
P
V
vnd
C
Nuclear DL Levels
nc12
nc13
nc14
-35 -15 0 10 25 85
Time (min)
nc14a
nc14b
nc14b/c
Dark
vndEEE-MS2 00:00
00:12
00:22
D
Nuclear DL Levels
nc12
nc13
nc14
0 25
Time (min)
Light 10 min
vndEEE-MS2 00:00
00:12
00:22
E
vndEEE-MS2 in dl-LEXY
Dark
Light 10 min
### of spots
0
50
100
150
0 5 10 15 20
Time (min)

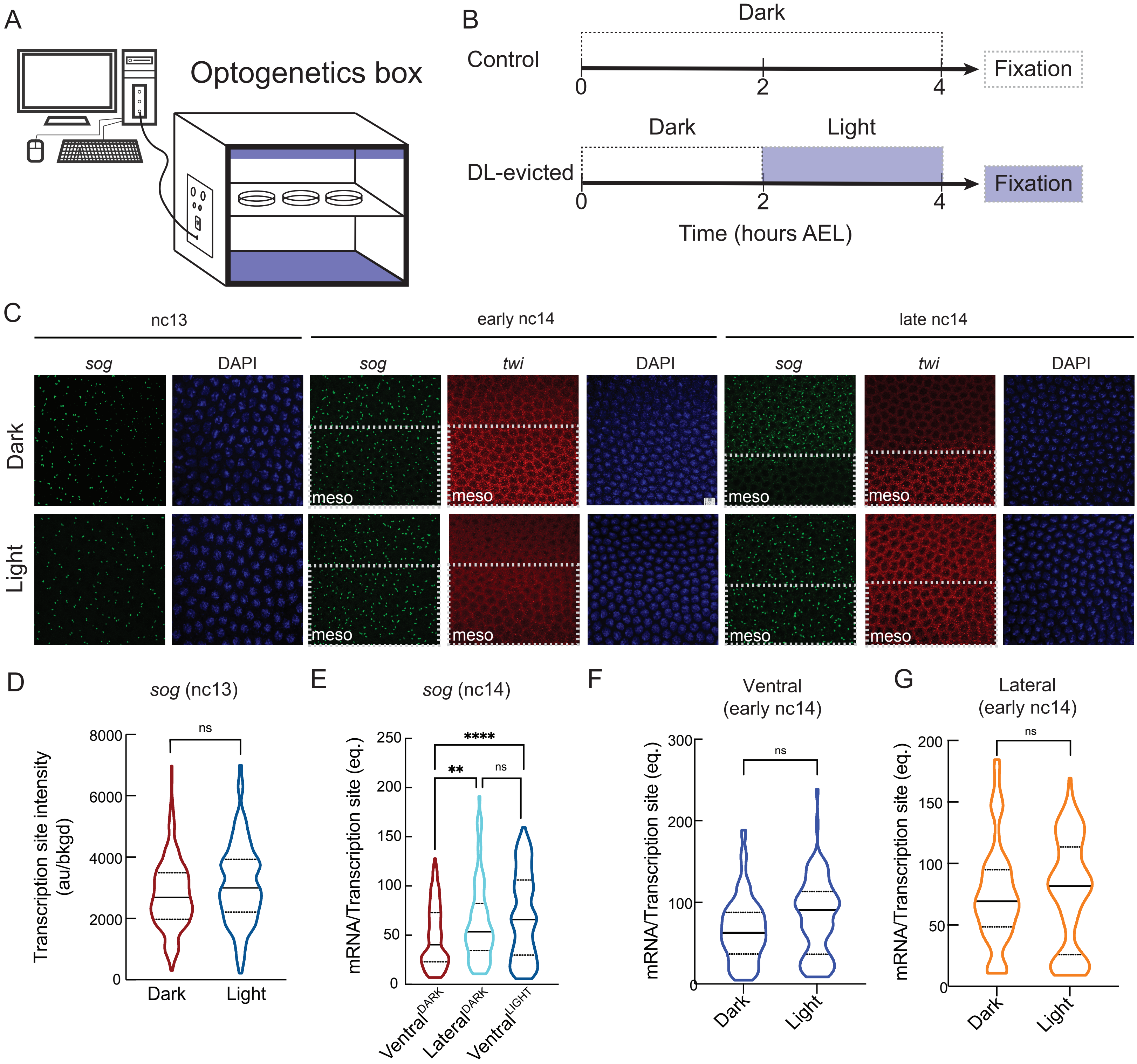
A
Optogenetics box
B
Control
Dark
Fixation
DL-evicted
Dark
Light
Fixation
0
2
4
Time (hours AEL)
C
nc13
early nc14
late nc14
sog
DAPI
sog
twi
DAPI
sog
twi
DAPI
Dark
Light
meso
D
sog (nc13)
Transcription site intensity (au/bkgd)
ns
Dark
Light
E
sog (nc14)
mRNA/Transcription site (eq.)
****
**
ns
Ventral^DARK
Lateral^DARK
Ventral^LIGHT
F
Ventral (early nc14)
mRNA/Transcription site (eq.)
ns
Dark
Light
G
Lateral (early nc14)
mRNA/Transcription site (eq.)
ns
Dark
Light

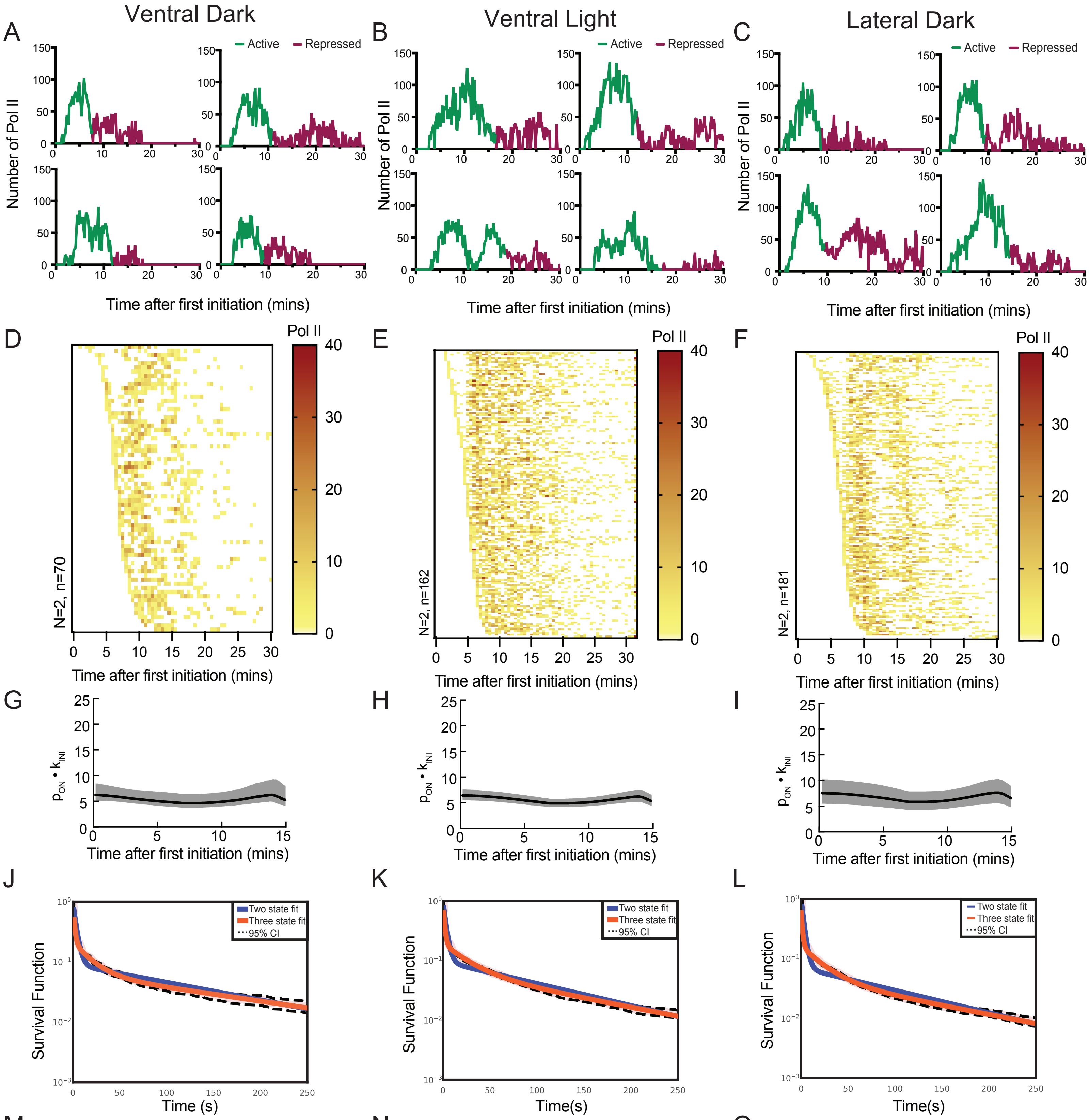


M

| | ON | OFF1 | OFF2 |
|---|---|---|---|
| Duration (s) | 7.76 | 179.14 | 15.19 |
| Min. | 7.76 | 179.14 | 15.19 |
| Max. | 7.76 | 179.14 | 15.19 |
| Probability | 0.12 | 0.72 | 0.16 |
| Min. | 0.12 | 0.72 | 0.16 |
| Max. | 0.12 | 0.72 | 0.16 |

N

| | ON | OFF1 | OFF2 |
|---|---|---|---|
| Duration (s) | 8.60 | 134.22 | 21.87 |
| Min. | 8.60 | 134.22 | 21.87 |
| Max. | 8.60 | 134.22 | 21.87 |
| Probability | 0.13 | 0.63 | 0.23 |
| Min. | 0.13 | 0.63 | 0.23 |
| Max. | 0.13 | 0.63 | 0.23 |

O

| | ON | OFF1 | OFF2 |
|---|---|---|---|
| Duration (s) | 8.86 | 136.38 | 19.23 |
| Min. | 8.86 | 136.38 | 19.23 |
| Max. | 8.86 | 136.38 | 19.23 |
| Probability | 0.15 | 0.60 | 0.25 |
| Min. | 0.15 | 0.60 | 0.25 |
| Max. | 0.15 | 0.60 | 0.25 |
